# Coordination of RNA and protein condensation by the P granule protein MEG-3

**DOI:** 10.1101/2020.10.15.340570

**Authors:** Helen Schmidt, Andrea Putnam, Dominique Rasoloson, Geraldine Seydoux

## Abstract

Germ granules are RNA-protein condensates in germ cells. The mechanisms that drive germ granule assembly are not fully understood. MEG-3 is an intrinsically-disordered protein required for germ (P) granule assembly in *C. elegans*. MEG-3 forms gel-like condensates on liquid condensates assembled by PGL proteins. MEG-3 is related to the GCNA family and contains an N-terminal disordered region (IDR) and a predicted ordered C-terminus featuring an HMG-like motif (HMGL). Using *in vitro* and *in vivo* experiments, we find the MEG-3 C-terminus is necessary and sufficient to build MEG-3/PGL co-condensates independent of RNA. The HMGL domain is required for high affinity MEG-3/PGL binding *in vitro* and for assembly of MEG-3/PGL co-condensates *in vivo*. The MEG-3 IDR binds RNA *in vitro* and is required but not sufficient to recruit RNA to P granules. Our findings suggest that P granule assembly depends in part on protein-protein interactions that drive condensation independent of RNA.

## INTRODUCTION

In animals with germ plasm, specification of the germline depends on the segregation of maternal RNAs and proteins (germline determinants) to the primordial germ cells. Germline determinants assemble in germ granules, micron-sized dense assemblies that concentrate RNA and RNA-binding proteins (Jamieson-Lucy and Mullins, 2019; Marnik and Updike, 2019; Seydoux, 2018; Trcek and Lehmann, 2019). Superficially, germ granules resemble RNA-rich condensates that form in the cytoplasm of somatic cells, including P bodies and stress granules. In recent years, much progress has been made in our understanding of stress granule assembly with the realization that stress granules resemble liquid condensates that assemble by liquid-liquid phase separation (LLPS). LLPS is a thermodynamic process that causes interacting molecules to dynamically partition between a dense condensed phase and a more dilute phase (e.g. the cytoplasm) (Banani et al., 2017; Mitrea and Kriwacki, 2016). Low-affinity binding interactions, often involving disordered and RNA-binding domains, are sufficient to drive LLPS of proteins and RNA in reconstituted systems (Lin et al., 2015; Molliex et al., 2015; Zagrovic et al., 2018). The ability of RNA to phase separate in the absence of proteins *in vitro* has also been proposed to contribute to RNA granule assembly *in vivo*, especially in the case of stress granules which arise under conditions of general translational arrest (Tauber et al., 2020; Van Treeck et al., 2018). An emerging model is that the combined action of many low-affinity interactions between RNA molecules and multivalent RNA-binding proteins create RNA-based protein networks that drive LLPS (Guillén-Boixet et al., 2020; Sanders et al., 2020; Yang et al., 2020; Zhang et al., 2015).

Unlike the dynamic condensates assembled by LLPS *in vitro*, germ granules are not well-mixed, single-phase liquid droplets. High resolution microscopy has revealed that germ granules are heterogenous assemblies of dynamic and less dynamic condensates that co-assemble but do not fully mix. For example, Drosophila germ granules contain non-dynamic RNA clusters embedded in dynamic, protein-rich condensates (Little et al., 2015; Niepielko et al., 2018; Trcek et al., 2015). Germ granules in zebrafish and Xenopus are built on an amyloid-like scaffold that organizes mRNAs in non-overlapping, transcript-specific zones (Boke et al., 2016; Fuentes et al., 2018; Roovers et al., 2018). The mechanisms that bring together condensates with different material properties and their contribution to RNA recruitment in germ granules are not well understood.

In this study, we examine the assembly of P granules, germ granule in *C. elegans*. At the core of P granules are liquid condensates assembled by PGL proteins. PGL-1 and PGL-3 are self-dimerizing, RGG domain proteins that readily form condensates able to recruit other P granule components, such as the VASA-related RNA helicase GLH-1 (Aoki et al., 2016; Hanazawa et al., 2011; Saha et al., 2016; Updike et al., 2011). PGL condensates exist in germ cells throughout oogenesis and are maternally-inherited by the embryo. In newly fertilized zygotes, the surface of PGL condensates becomes covered by smaller condensates assembled by MEG-3 and MEG-4, two homologous intrinsically-disordered proteins (Wang et al., 2014). Unlike PGL condensates, MEG-3 condensates resist dilution and salt challenge, consistent with a gel-like material (Putnam et al., 2019). (In this study, we use the term condensate to refer to concentrated protein assemblies that self-assemble without implying a mechanism for assembly, which could involve aggregation, LLPS or other mechanisms). During zygote polarization, MEG-3 and MEG-4 condensates enrich with other germ plasm components in the posterior cytoplasm (Putnam et al., 2019; Smith et al., 2016; Wang et al., 2014). This relocalization correlates with preferential growth of MEG-coated PGL droplets in the posterior and dissolution of “naked” PGL droplets in the anterior side (Brangwynne et al., 2009; Smith et al., 2016). In addition to PGL and MEG co-assemblies, P granules also concentrate specific maternal transcripts (Parker et al., 2020; Seydoux and Fire, 1994). A survey of mRNAs that immunoprecipate with PGL-1 and MEG-3 suggest that MEG-3 is most directly responsible for recruiting mRNAs to P granules (Lee et al., 2020). MEG-3 binds to ~500s maternal mRNAs, including transcripts coding for germline determinants. Recruitment of mRNAs to P granules ensures their preferential segregation to the primordial germ cells. Embryos lacking MEG-3 and MEG-4 do not localize PGL droplets, do not condense P granule-associated mRNAs, and display partially penetrant (30%) sterility (Lee et al., 2020; Wang et al., 2014)

To understand how MEG-3 coordinates PGL and RNA condensation, we used *in vitro* reconstitution experiments and genome editing of the *meg-3* locus to define functional domains in MEG-3. We find that MEG-3 is a bifunctional protein with separate domains for RNA recruitment and protein condensation. We identify a predicted ordered motif in the MEG-3 C-terminus required for high affinity binding to PGL-3 *in vitro* and find that this domain is essential to build MEG-3/PGL-3 coassemblies that recruit RNA *in vivo*. The MEG-3_IDR_ binds RNA and enriches MEG-3 in germ plasm but is not sufficient on its own to assemble RNA-rich condensates. Our observations highlight the importance of condensation driven by proteinprotein interactions in the assembly of germ granules.

## RESULTS

### The MEG-3 IDR and C-terminus synergize to promote MEG-3 condensation *in vitro*

IUPred2A (Mészáros et al., 2018) predicts in the MEG-3 sequence a 544 residue N-terminal domain with high disorder (MEG-3_IDR_, aa1-544) and a 318 residue C-terminal domain (MEG-3_Cterm_, aa545-862) with lower disorder (Figure 1A, B). MEG-3_Cterm_ contains a region (aa700-744) with sequence similarity to the HMG-like motif found in the GCNA family of intrinsically-disordered proteins (Figure 1C). GCNA family members also contain long N-terminal disordered domains, but these do not share sequence homology with the MEG-3 IDR (Carmell et al., 2016).

**Figure 1:**
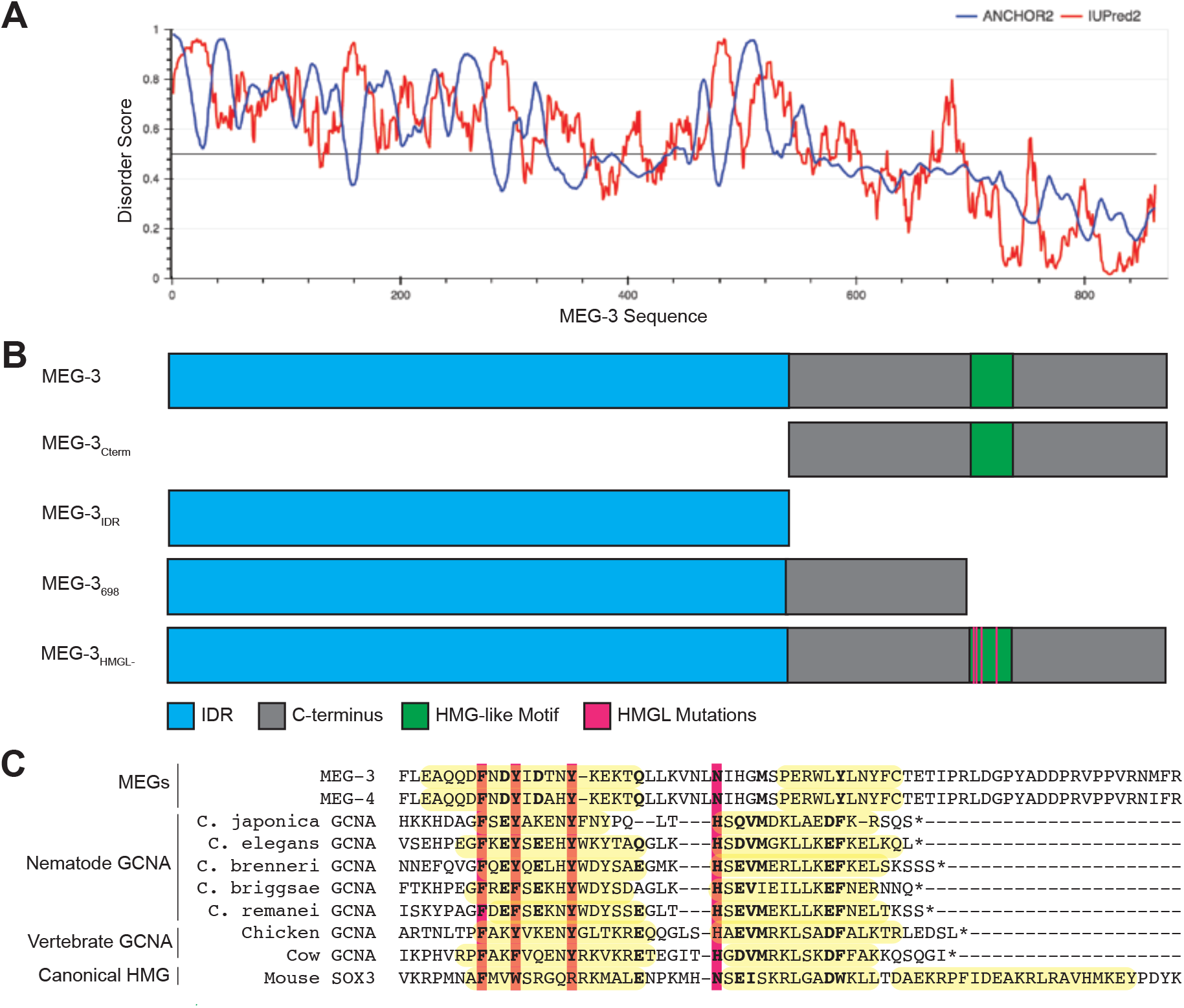
Domain organization of MEG-3. (**A**) MEG-3 amino-acid sequence (N to C-terminus) on the X-axis is plotted against disorder score on the Y-axis as predicted by ANCHOR2 (blue) and IUPred2 (red) (Mészáros et al., 2018) with a range from zero to one, where one is the most disordered. (**B**) Schematics of wild-type MEG-3 and four MEG-3 variants analyzed in this study. Amino acid positions are aligned with A. The disordered region (blue), C-terminus (grey), and HMG-like motif (green) are indicated. Magenta bars (alanine substitutions) correspond to four conserved residues in the HMG-like motif shaded in magenta in C. (**C**) Alignment of the HMG-like motif in MEG-3 and MEG-4 with the HMG-like motif in GCNA proteins (Carmell et al., 2016) and the canonical HMG box of mouse SOX3. Amino acids predicted to form alpha-helices are highlighted in yellow (Drozdetskiy et al., 2015). Bold indicates positions with greater than 70% amino acid similarity. Magenta bars indicate residues mutated to alanine in MEG-3_HMGL-_.

To determine which regions of MEG-3 are required for condensation *in vitro*, we expressed and purified His-tagged full-length MEG-3 and four derivatives: MEG-3_Cterm_, MEG-3_IDR_, MEG-3_698_ an extended version of MEG-3_IDR_ terminating right before the HMG-like motif, and MEG-3_HMGL-_, a full length MEG-3 variant with alanine substitutions in 4 conserved residues in the HMG-like motif (Figure 1B, C). The proteins were trace-labeled with covalently attached fluorophores and condensation was assayed as a function of protein concentration in the presence of 150mM NaCl and 20ng/μl of *nos-2* RNA (*nos-2* is an mRNA found in P granules (Lee et al., 2020; Subramaniam and Seydoux, 1999, p. 1); Methods). Wild-type MEG-3 forms small condensates at 50nM, a concentration similar to that estimated for MEG-3 *in vivo* (Putnam et al., 2019; Saha et al., 2016). With increasing protein concentration, the fraction of MEG-3 in condensates increases (Figure 2A and B). MEG-3_HMGL-_ behaved indistinguishably from wild-type. In contrast, the MEG-3_Cterm_ and MEG-3_IDR_ lagged behind, with the MEG-3_IDR_ lagging the most at the lowest concentration (Figure 2B). MEG-3_698_ behaved like MEG-3_IDR_ (Figure 2 – figure supplement 1A). We conclude that both the IDR and C-terminus contribute to MEG-3 condensation *in vitro*.

**Figure 2:**
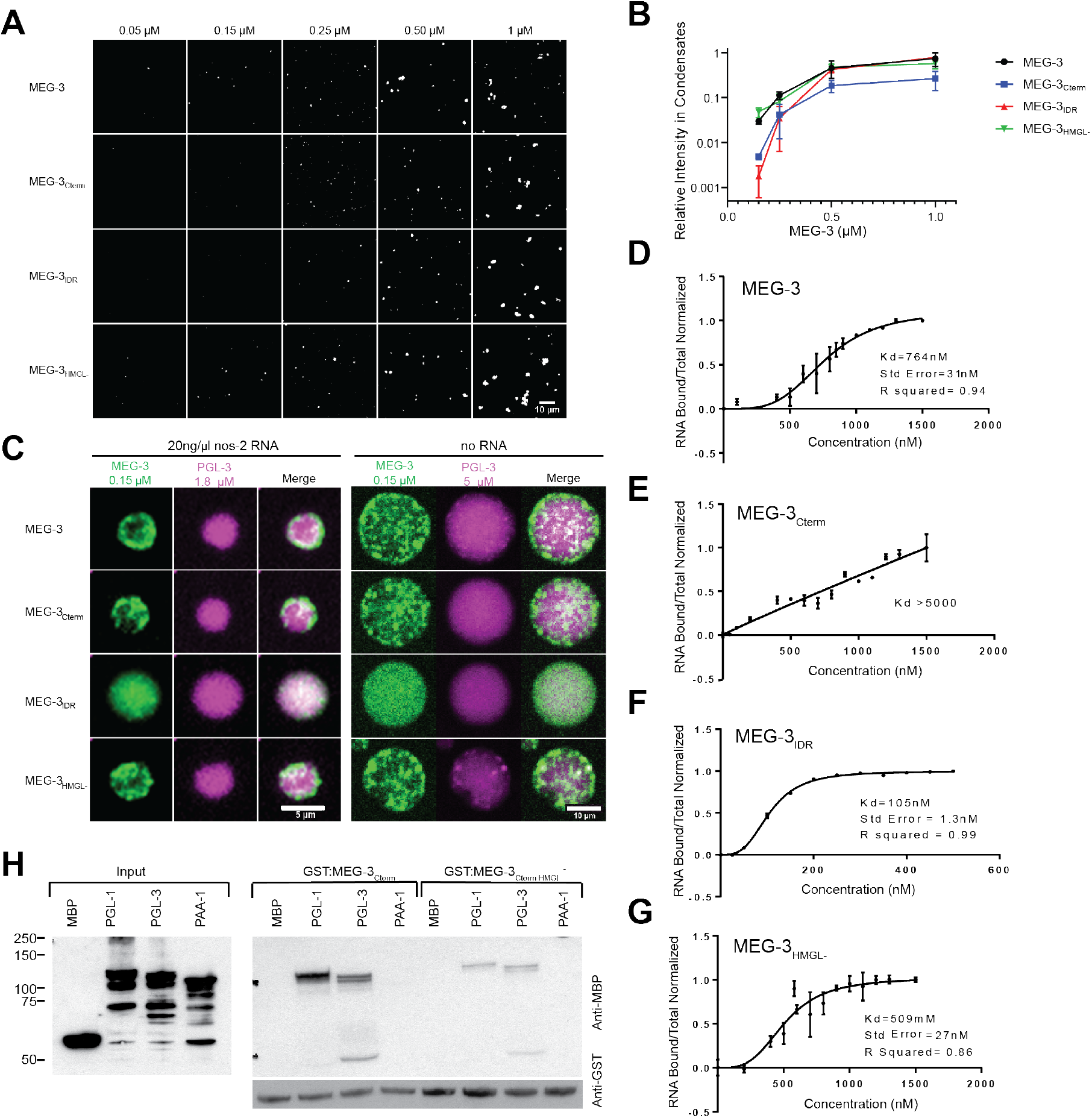
in vitro characterization of wild-type MEG-3 and variants. **(A)** Representative photomicrographs of Alexa647 trace-labeled MEG-3 and MEG-3 variants, at increasing concentration from left to right, in condensation buffer with 20 ng/μL *nos-2* mRNA. Quantification of MEG-3 in condensates is shown in B. **(B)** Total intensity of MEG-3 in condensates (Y axis, normalized to 1 μM full length MEG-3) plotted against MEG-3 concentration (X axis). Condensates were assembled as in A. Dots indicate the mean of 3 replicates and errors bars the standard deviation. **(C)** Representative photomicrographs of MEG-3 and MEG-3 derivatives (trace labeled with Alexa647) incubated for 30 mins with PGL-3 (trace labeled with Alexa488) with and without nos-2 mRNA in condensation buffer. **(D-G)** RNA binding curves for MEG-3 (D), MEG-3_Cterm_ (E), MEG-3_IDR_ (F), and MEG-3_HMGL-_ (G). Protein concentration is plotted on the X-axis. The ratio of bound RNA to total RNA, normalized to the ratio at the maximum concentration is plotted on the Y-axis. Dots indicate the mean of four replicates and the error bars the standard deviation (not shown when <0.06). Curve fit and Kd calculation based on specific binding with Hill slope (Methods). **(H)** Analysis of GST::MEG-3_Cterm_ and MBP::PGL-1 and MBP::PGL-3 interactions by GST-pull down assay with MBP and MBP::PAA-1 as negative controls. Western blots of E. coli lysates expressing the indicated MBP-fusions before (Input) and after immobilization on magnetic beads with the indicated GST-fusions. Western blot of GST-fusions is shown below. See Figure 2 – figure supplement 1C for quantification of additional replicates.

### Co-assembly of MEG-3/PGL-3 condensates *in vitro* is driven by the MEG-3 C-terminus and does not require RNA or the MEG-3 IDR

When combined in condensation assays, MEG-3 and PGL-3 form co-condensates that resemble the architecture of P granules *in vivo*, with the smaller MEG-3 condensates (~100 nm) forming a dense layer on the surface of the larger PGL-3 condensates (Putnam et al., 2019). MEG-3_Cterm_ and MEG-3_HMGL-_ formed co-condensates with PGL-3 that were indistinguishable from those formed by wild-type MEG-3(Figure 2C). The MEG-3_IDR_, in contrast, failed to assemble condensates on the surface of PGL-3, or away from PGL-3, and instead mixed homogenously with the PGL-3 phase as previously reported (Putnam et al. 2019, Figure 2C).

We repeated the co-condensation assays in the absence of RNA using a higher concentration of PGL to force PGL condensation in the absence of RNA. MEG/PGL co-condensates assembled under those conditions were indistinguishable from co-condensates assembled in the presence of RNA (Figure 2C). Again, the C-terminus was necessary and sufficient for co-assembly. MEG-3_IDR_ homogenously mixed with the PGL-3 phase and did not form independent condensates, confirming that MEG-3_IDR_ is solubilized by PGL-3. We conclude that the MEG-3 C-terminus is the primary driver of MEG-3 condensation and that condensation of MEG-3 on PGL condensates depends on the MEG-3 C-terminus and does not require RNA *in vitro*.

### The MEG-3 IDR is necessary and sufficient for RNA binding *in vitro*

Using fluorescence polarization and gel shift assays, we previously showed that the MEG-3_IDR_ binds an RNA oligo (poly-U30) with near nanomolar affinity *in vitro* (Smith et al., 2016). We repeated these observations using a filter binding assay where proteins are immobilized on a filter to minimize possible interference due to condensation of MEG-3 in solution (Methods, Figure 2D-G). Consistent with previous observations (Smith et al., 2016), we found that the MEG-3_IDR_ exhibits high affinity for RNA (Kd=105 nM; Figure 2F). MEG-3_698_ also exhibited high affinity (Kd=95 nM; Figure 2 – figure supplement 1B). Wild-type MEG-3 bound RNA efficiently (Figure 2D), albeit at a lower affinity than MEG-3_IDR_ and MEG-3_698_. In contrast, MEG-3_Cterm_ exhibits negligible RNA binding (Figure 2E). HMG domains are common in DNA-binding proteins and have been shown to mediate protein:nucleic acid interactions *in vivo* (Genzor and Bortvin, 2015; Reeves, 2001; Thapar, 2015), raising the possibility that the HMG-like domain in MEG-3 might contribute to RNA binding. We found, however, that the MEG-3_HMGL-_ bound to RNA with high affinity, similar to wild-type MEG-3 (Figure 2G). We conclude that the HMG-like domain does not contribute to RNA binding, which is driven primarily by the IDR.

### The HMG domain is required for high affinity binding to PGL proteins *in vitro*

HMG domains have also been implicated in protein-protein interactions (Reeves, 2001; Stros et al., 2007; Wilson and Koopman, 2002). We reported previously that MEG-3 binds directly to PGL-1, as determined in a GST-pull down assay using partially purified recombinant proteins (Wang et al., 2014). We repeated this assay using fusion proteins of GST::MEG-3_Cterm_ and MBP::PGL-1 and PGL-3. GST::MEG-3_IDR_ fusions were not expressed and thus could not be tested in this assay. We found that the GST:MEG-3_Cterm_ binds efficiently to PGL-1 and PGL-3, but not to MBP or to an unrelated control protein PAA-1. Remarkably, we found that the HMG-like motif contributes to these interactions. A GST::MEG-3_Cterm_ fusion with mutations in the HMG-like domain bound less efficiently to PGl-1 and PGL-3 (Figure 2H, Figure 2 – figure supplement 1C). We conclude that the MEG-3_Cterm_ is sufficient to bind to PGL proteins *in vitro* and that these interactions require the HMG-like motif for high efficiency binding.

### The MEG-3 C-terminus is the primary driver of MEG-3 condensation *in vivo*

To test the functionality of MEG-3 domains *in vivo*, we used CRISPR genome editing to re-create at the *meg-3* locus the same variants analyzed *in vitro*. We used a *C. elegans* strain line with the *meg-4* locus deleted to avoid possible complementation by MEG-4. To allow visualization of MEG-3 protein by immunofluorescence, each variant (and wild-type *meg-3*) was tagged with a C-terminal OLLAS peptide. We avoided the use of fluorescent tags as fluorescent tags have been reported to affect the behavior of proteins in P granules (Uebel and Phillips, 2019).

As reported previously for untagged MEG-3 (Wang et al., 2014), MEG-3 tagged with OLLAS could be detected diffusively in the cytoplasm and in condensates. Before polarization, MEG-3 was uniformly distributed throughout the zygote. After polarization, MEG-3 in the cytoplasm and in condensates became enriched in the posterior half of the zygote destined for the germline blastomere P_1_ (“germ plasm”). MEG-3 continued to segregate preferentially with P blastomeres in subsequent divisions (P_1_ through P_4_) (Figure 3A).

**Figure 3:**
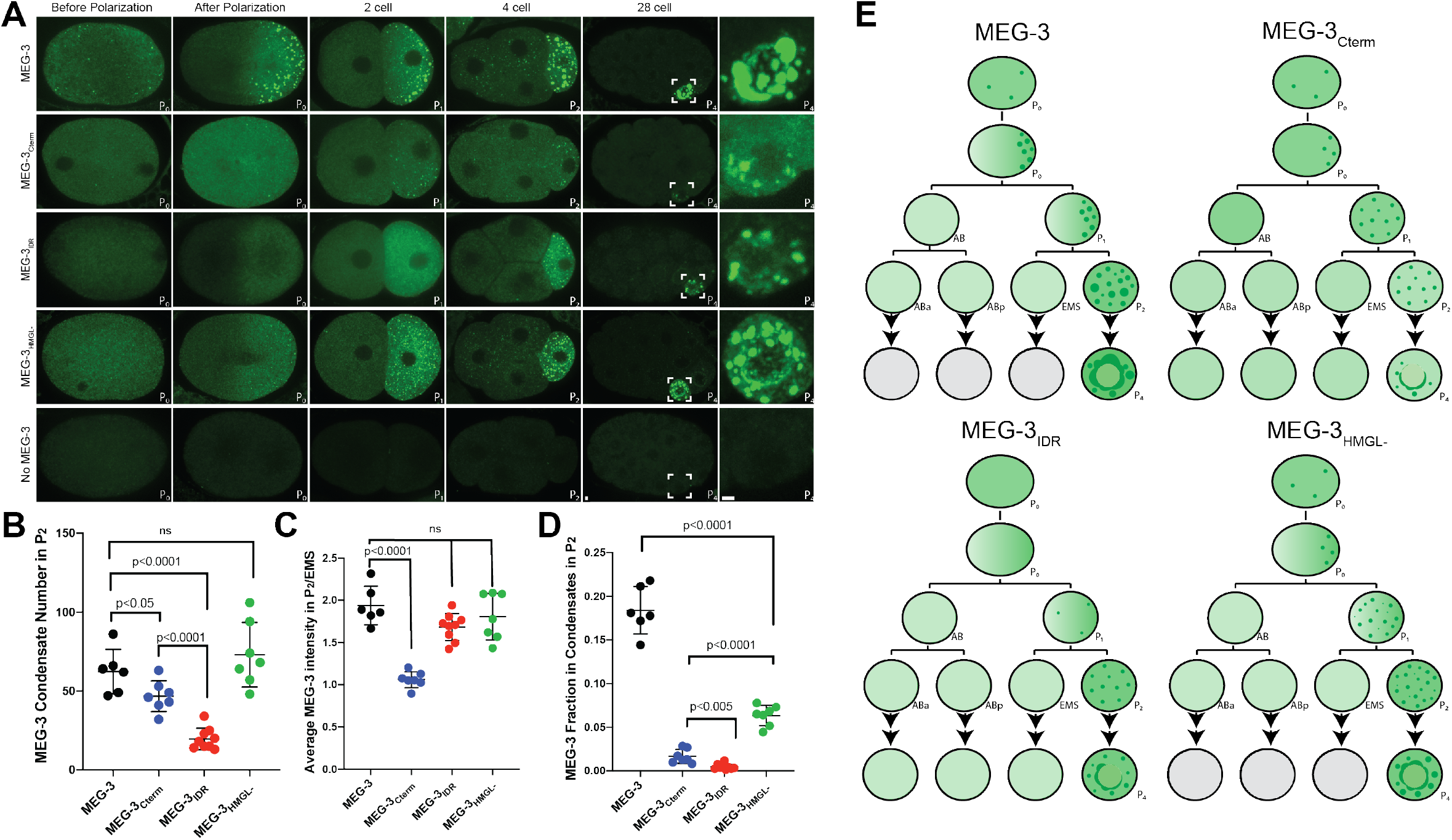
Localization of wild-type MEG-3 and variants in early embryos. **(A)** Representative photomicrographs of embryos immunostained for OLLAS and expressing the indicated OLLAS-tagged MEG-3 derivatives. Last row shows *meg-3 meg-4* embryos as negative control for OLLAS staining. Images are representative of each stage as indicated (Before and after polarization are 1-cell stage zygotes). A minimum of 3 embryos from two independent experiments were analyzed for each stage. Scale bars are 1μm. All images are normalized to same fluorescent intensity range except for the last column showing high magnification views of P_4_ from 28-cell stage image adjusted to highlight MEG-3 granules. **(B)** Scatterplot showing the number of MEG-3 condensates in P_2_ in embryos expressing the indicated MEG-3 derivatives. Each dot represents an embryo. **(C)** Scatterplot showing enrichment of MEG-3 in P_2_ over the somatic blastomere EMS, calculated by dividing the average intensity in P_2_ by the average intensity in EMS. Each dot represents an embryo also included in the analysis shown in B. **(D)** Scatterplot showing the fraction the MEG-3 signal localized to condensates over total signal in P_2_. Each dot represents an embryo also included in the analysis in B. **(E)** Summary of MEG-3 (green) distribution derived from data presented in A. Horizontal lines and arrows denote one cell division. Note that wild-type MEG-3 and MEG-3_HMGL-_ are rapidly turned over in somatic cells after the 4-cell stage (grey cells) as shown in Figure 3 – figure supplement 1C.

All four MEG-3 variants exhibited unique localization patterns distinct from wild-type. MEG-3_IDR_ enriched in posterior cytoplasm and segregated preferentially to P blastomeres but did not appear robustly in condensates until the 4-cell stage (P_2_ blastomere, Figure 3A). MEG-3_698_ behaved similarly to MEG-3_IDR_ (Figure 3 – figure supplement 1A). MEG-3_Cterm_ did not enrich asymmetrically in the cytoplasm but formed condensates in the zygote posterior and continued to form condensates only in P blastomeres despite being present in the cytoplasm of all cells (Figure 3A). MEG-3_HMGL-_ behaved most similarly to wild-type MEG-3 enriching in the zygote posterior and forming condensates as early as the 1-cell stage, although the condensates appeared smaller at all stages (Figure 3A).

For each MEG-3 derivative, we quantified the number of condensates and the degree of enrichment in the P blastomere over somatic blastomeres and in condensates over the cytoplasm. Wild-type MEG-3 and MEG-3_HMGL-_ formed a similar number of condensates, while MEG-3_Cterm_ formed fewer and MEG-3_IDR_ the least in the 4-cell stage (Figure 3B). The MEG-3_Cterm_ did not enrich in the P_2_ blastomere, whereas MEG-3_IDR_ and MEG-3_HMGL-_ enriched as efficiently as wild-type (Figure 3C). Finally, none of MEG-3 derivatives enriched in condensates as efficiently as wildtype (Figure 3D).

After the four-cell stage, the low levels of wild-type MEG-3 and MEG-3_HMGL-_ inherited by somatic blastomeres were rapidly cleared. In contrast, MEG-3_IDR_ and MEG-3_Cterm_ persisted in somatic blastomeres at least until the 28-cell stage (Figure 3 – figure supplement 1B). Western analyses revealed that MEG-3 and MEG-3_HMGL-_ accumulate to similar levels, whereas MEG-3_IDR_ and MEG-3_Cterm_ were more abundant in mixed-stage embryo lysates, consistent with slower turnover in somatic lineages (Figure 3 – figure supplement 1C).

The condensation, segregation, and turnover patterns of MEG-3, MEG-3_IDR_, MEG-3_Cterm_ MEG-3_HMGL-_ are summarized in Figure 3E. From this analysis, we conclude that: 1) the MEG-3_IDR_ is necessary and sufficient for enrichment of cytoplasmic MEG-3 in germ plasm, 2) the MEG-3 C-terminus is necessary and sufficient for condensation of MEG-3 in germ plasm starting in the zygote stage, 3) the HMG-like motif is required for efficient MEG-3 condensation, and 4) both the C-terminus and the IDR are required for timely turn-over of MEG-3 in somatic lineages.

### Co-assembly of MEG-3/PGL-3 condensates *in vivo* is driven by the MEG-3 C-terminus and requires the HMGL motif

MEG-3 and MEG-4 are required redundantly to localize PGL condensates to the posterior of the zygote for preferential segregation to the P lineage (Smith et al., 2016; Wang et al., 2014). To examine the distribution of PGL condensates relative to MEG-3 condensates, we utilized the KT3 and OLLAS antibodies for immunostaining of untagged endogenous PGL-3 and OLLAS-tagged MEG-3. In embryos expressing wild-type MEG-3, MEG-3 and PGL-3 co-localize in posterior condensates that are segregated to the P_1_ blastomere. (Figure 4A). In embryos lacking *meg-3* and *meg-4*, PGL-3 condensates distributed throughout the cytoplasm of the zygote and segregated equally to AB and P_1_ (Figure 4A). We observed a similar pattern in embryos expressing MEG-3_IDR_, MEG-3_698_ and MEG-3_HMGL-_ indicating that none of these MEG-3 derivatives are sufficient to localize PGL condensates (Figure 4A, Figure 3 – figure supplement 1A). In contrast, in embryos expressing MEG-3_Cterm_, PGL-3 condensates localized properly in P_1_, although they were smaller and fewer than in wild-type (Figure 4A and B). Embryos expressing MEG-3_Cterm_ enriches PGL-3 in P_1_, though not as efficiently as wild-type, while PGL-3 is not enriched in *meg-3 meg-4*, or embryos expressing MEG-3_IDR_ or MEG-3_HMGL-_ (Figure 4C). In wild-type 28-cell stage embryos, PGL-3 condensates are highly enriched in P_4_. No such enrichment was observed in embryos expressing the MEG-3_Cterm_ or any other MEG-3 variant (Figure 4 – figure supplement 1). We conclude that the MEG-3_Cterm_ is sufficient to enrich PGL-3 condensates in P blastomeres in early stages, but not sufficient to support robust PGL-3 localization through P_4_.

**Figure 4:**
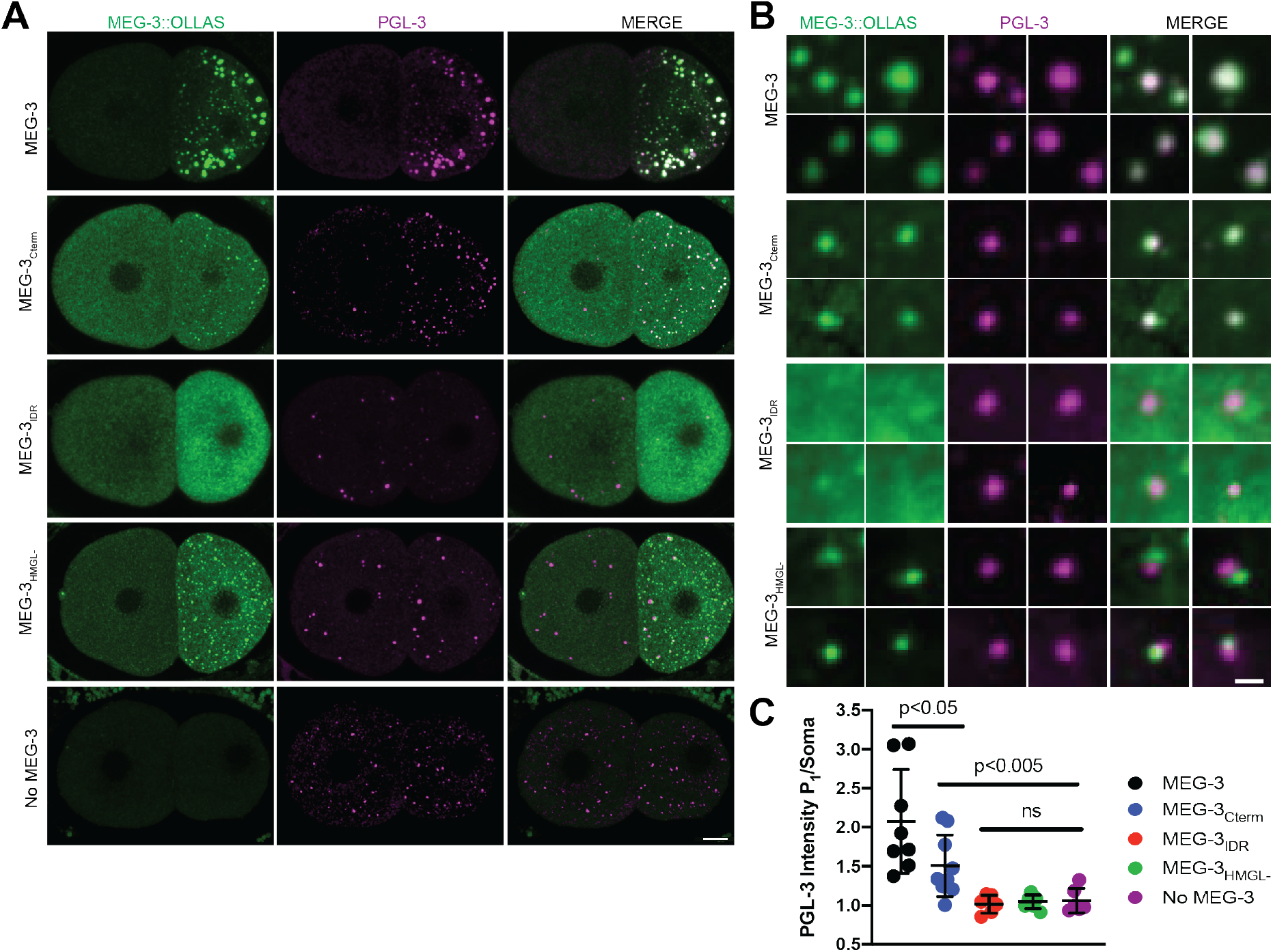
Localization of PGL-3 relative to wild-type MEG-3 and variants. **(A)** Representative photomicrographs of two-cell embryos expressing the indicated MEG-3 mutants and immunostained for MEG-3 (anti-OLLAS antibody) and PGL-3 (anti-PGL-3 antibody). Scale bar is 5 μm. **(B)** High magnification photomicrographs of individual MEG-3/PGL-3 assemblies in embryos expressing the indicated MEG-3 derivatives. White color in the merge indicates overlap. Scale bar is 1μm. **(C)** Scatterplot of the enrichment of PGL-3 in P_1_ calculated by dividing the average intensity in P_1_ by the average intensity in AB. Each dot represents an embryo.

Wild-type MEG-3 condensates associate closely with the surface of PGL condensates (Putnam et al., 2019; Wang et al., 2014). With the resolution afforded by immunostaining, this configuration appears as co-localized MEG and PGL puncta in fixed embryos (Wang et al., 2014, Figure 4B). We found that PGL-3 condensates co-localized with MEG-3_Cterm_ condensates (37/37 PGL-3 condensates scored in P_1_; Figure 4B) as in wild-type. In contrast, we observed no such colocalization with MEG-3_IDR_ or MEG-3_HMGL-_. The MEG-3_IDR_ is mostly cytoplasmic and forms only rare condensates in P_2_. We occasionally observed PGL condensates with an adjacent MEG-3_IDR_ condensate (5/19 PGL-3 condensates scored in P_1_, Figure 4B), but these were not co-localized. Unlike the MEG-3_IDR_, MEG-3_HMGL-_ forms many condensates in P_2_, although these tended to be smaller than wild-type (Figure 3A,B). Still, although we occasionally observed PGL condensates with an adjacent MEG-3_HMGL-_ condensate (12/30 PGL-3 condensates scored in P_1_; Figure 4B), we never observed fully overlapping PGL/MEG-3_HMGL-_ co-condensates. We conclude that, despite forming many condensates in P blastomeres, MEG-3_HMGL-_ condensates do not associate efficiently with, and do not support the localization of, PGL-3 condensates.

### Efficient recruitment of Y51F10.2 mRNA to P granules requires the MEG-3 IDR, C-terminus and HMG-like motif

MEG-3 recruits mRNAs to P granules by direct binding which traps mRNA into the non-dynamic MEG-3 condensates (Lee et al., 2020). To determine which MEG-3 domain is required for mRNA recruitment to MEG-3 condensates *in vivo*, we performed *in situ* hybridization against the MEG-3-bound mRNA *Y51F10.2*. Prior to polarization, *Y51F10.2* is uniformly distributed throughout the zygote cytoplasm (Figure 5A). *Y51F10.2* becomes progressively enriched in P granules starting in the late 1-cell stage and forms easily detectable micron-sized foci by the 4-cell stage ((Lee et al., 2020), Figure 5A). In contrast, in *meg-3 meg-4* embryos, *Y51F10.2* remains uniformly distributed in the cytoplasm at all stages. Strikingly, we observed the same failure to assemble *Y51F10.2* foci in embryos expressing MEG-3_IDR_, MEG-3_Cterm_ and MEG-3_HMGL-_ (Figure 5A). This was surprising since all three MEG-3 variants form visible condensates by the 4-cell stage (Figure 3A). We note that lack of PGL enrichment in posterior cytoplasm is unlikely to cause this RNA defect since *pgl-1;pgl-3* mutants still assemble *Y51F10.2* clusters (Lee et al., 2020).

**Figure 5:**
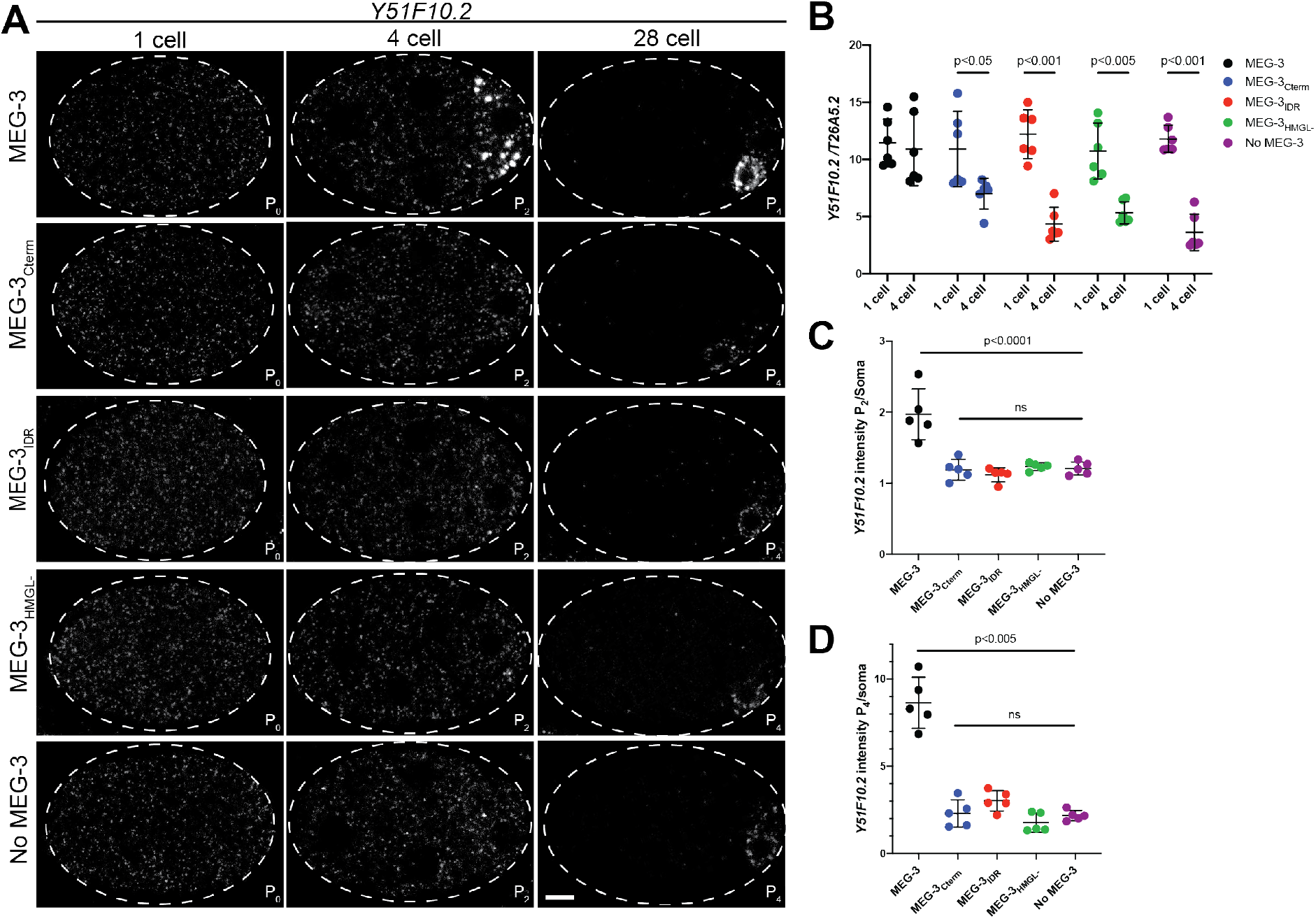
Distribution of *Y51F10.2* mRNA in embryos expressing wild-type MEG-3 and variants. **(A)** Representative photomicrographs of single confocal slices of fixed embryos expressing the indicated MEG-3 variants and hybridized to fluorescent probes complementary to the P granule-enriched mRNA *Y51F10.2* (white signal). **(B)** Scatterplot showing the ratio of *Y51F10.2* to *T26A5.2* mRNA signal in P_0_ and P_2_ embryos expressing the indicated MEG-3 derivatives. Each dot represents an embryo. See Supplementary Figure 5 for *T26A5.2* mRNA localization and levels **(C)** Scatterplot showing enrichment of *Y51F10.2* mRNA in P_2_ relative to somatic blastomeres in embryos expressing the indicated MEG-3 derivatives. Each dot represents an embryo (Methods). **(D)** Scatterplot showing enrichment of *Y51F10.2* mRNA in P_4_ relative to somatic blastomeres in embryos expressing the indicated MEG-3 derivatives. Each dot represents an embryo (Methods).

To characterize the fate of *Y51F10.2* transcripts in *meg-3meg-4* mutants, we compared the intensity of the *Y51F10.2 in situ* hybridization signal relative to a control RNA in 1-cell and 4-cell stage embryos (Methods, Figure 5 – figure supplement 1A,B). In wild-type, *Y51F10.2* RNA levels do not change significantly from the 1-cell to the 4-cell stage. In contrast, in *meg-3meg-4* embryos, *Y51F10.2* levels decreased by ~50% by the 4-cell stage, despite starting at levels similar to wild-type in the 1-cell stage. This finding is consistent with RNAseq results, which indicated lower levels of P granule mRNAs in *meg-3meg-4* embryos (Lee et al., 2020). We observed a similar loss of *Y51F10.2* RNA in embryos expressing MEG-3_IDR_, MEG-3_Cterm_ and MEG-3_HMGL-_ (Figure 5B). These results suggest that failure to recruit Y51F10.2 in granules leads to premature degradation.

After the 4-cell stage, as has been reported for other maternal RNAs (Baugh et al., 2003; Seydoux and Fire, 1994), *Y51F10.2* is rapidly turned over in somatic blastomeres. At the four cell stage, *Y51F10.2* mRNA levels are ~2-fold higher P_2_ than in somatic blastomeres in wild-type embryos, and ~1.2 fold higher in *meg-3 meg-4* embryos and in embryos expressing MEG-3_IDR_, MEG-3_Cterm_ and MEG-3_HMGL-_ (Figure 5C). By the 28-cell stage, in wild-type embryos, *Y51F10.2* levels are ~10-fold higher in the germline founder cell P_4_ compared to somatic blastomeres. In contrast, in *meg-3 meg-4* embryos, *Y51F10.2* mRNA levels were only ~2-fold enriched over somatic levels. Similarly, in embryos expressing the MEG-3_IDR_, MEG-3_Cterm_ and MEG-3_HMGL-_, *Y51F10.2* enrichment in P_4_ averaged around ~2-fold (Figure 5A, D).

Enrichment of mRNAs in P granules can also be detected using an oligo-dT probe to detect polyadenylated mRNAs (Seydoux and Fire, 1994). In wild-type 28cell stage embryos, strong poly-A signal is detected around the nucleus of the P_4_ blastomere (Figure 5 – figure supplement 1C). This perinuclear signal was absent in *meg-3 meg-4* mutants as well as in embryos expressing MEG-3_IDR_, MEG-3_Cterm_ and MEG-3_HMGL-_. The lack of polyA signal was particularly striking in the case of MEG-3_IDR_ and MEG-3_HMGL-_ since those variants assemble robust perinuclear condensates at this stage (Figure 3A).

Failure to efficiently segregate and stabilize maternal mRNAs in P blastomeres has been linked to the partial penetrance maternal effect sterility (~30%) of *meg-3meg-4* mutants (Lee et al., 2020). We observed similar levels of sterility in hermaphrodites derived from mothers expressing the MEG-3_IDR_, MEG-3_Cterm_ and MEG-3_HMGL-_ (Figure 5 – figure supplement 1D). We conclude that the MEG-3 c-terminus, IDR and HMG-like motif are all required for efficient mRNA recruitment to P granules, which in turn is required for enrichment and stabilization in the P lineage and robust germ cell fate specification.

## DISCUSSION

In this study, we have examined the function of the MEG-3 IDR and C-terminus in P granule assembly using recombinant proteins *in vitro* and genome editing *in vivo*. Our main findings (summarized in Figure 6) suggest the following model for germ granule assembly: the MEG-3 c-terminus including the HMGL domain mediates MEG:PGL protein interactions that stimulate MEG-3 condensation on the surface of PGL condensates. The MEG-3 IDR recruits RNA and amplifies MEG-3 condensation. MEG-3 condensation in turn stabilizes PGL condensates in the posterior and protects mRNAs from degradation ensuring their efficient segregation to the germline founder cell P_4_. Our findings suggest that germ granule assembly depends at least in part on protein-protein interactions that drive protein condensation independent of RNA.

**Figure 6:**
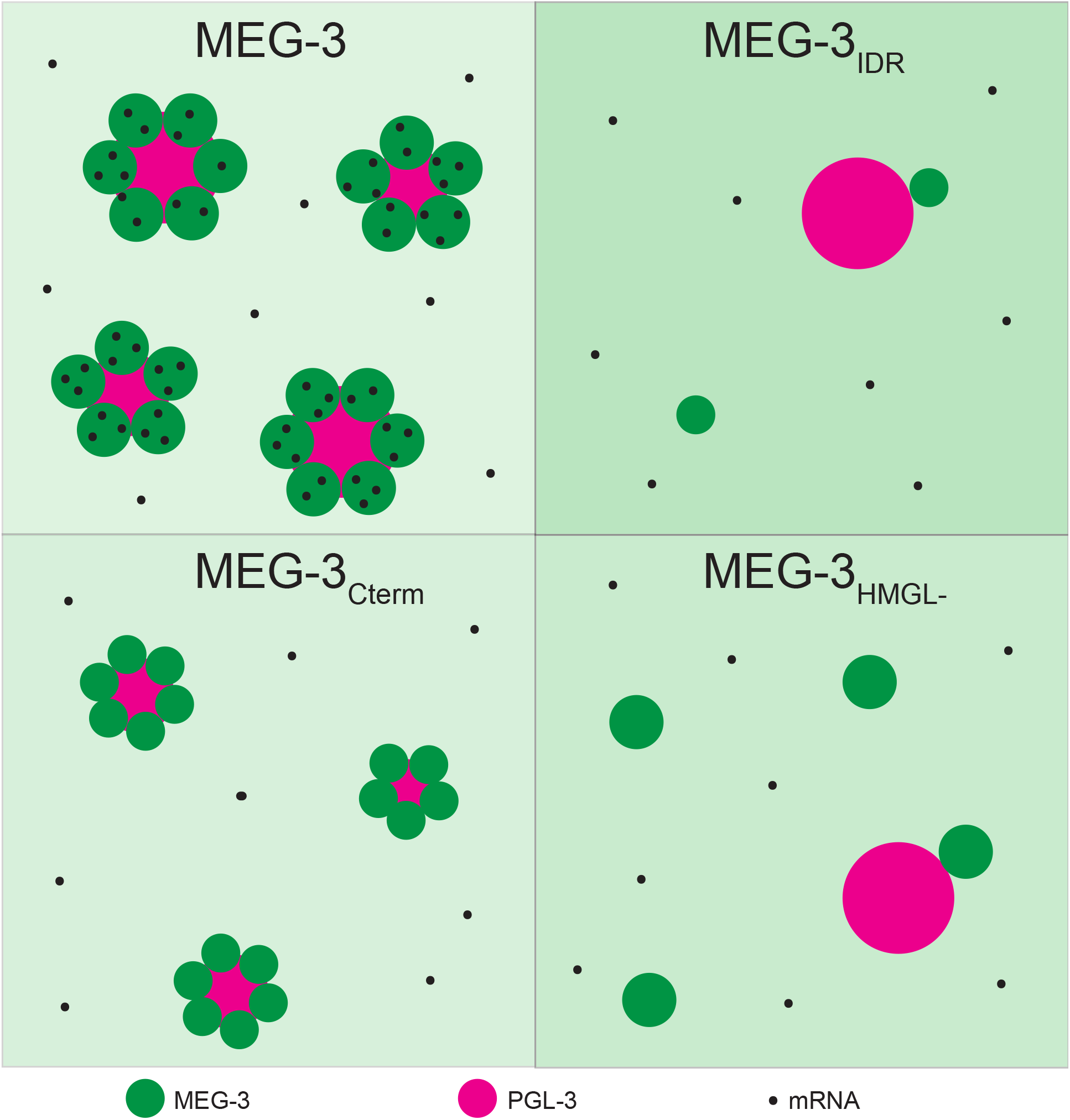
Diagram summarizing the distributions of MEG-3 and PGL-3 condensates and *Y51F10.2* mRNA in P blastomeres. In wild-type, MEG-3 is recruited to the surface of PGL droplets by its C-terminus while the MEG-3_IDR_ enriches RNA to the condensates. The MEG-3 does not interact with PGL efficiently and forms fewer condensates that do not enrich RNA. The MEG-3_Cterm_ is recruited to the surface of PGL droplets but, lacking the IDR, cannot enrich RNA. MEG-3_HMGL-_ does not interact with PGL efficiently and forms fewer condensates that do not enrich RNA.

### Similarities and differences between *in vitro* and *in vivo* observations

*In vitro* condensation assays using purified proteins and RNAs are powerful tools to identify domains and molecular interactions that drive condensate self-assembly (Li et al., 2018; Lin et al., 2015; Saha et al., 2016). Because these assays do not reconstitute the cytoplasmic environment, however, results need to be interpreted with caution. In this study, we compared the *in vitro* and *in vivo* behavior of wild-type MEG-3 and derivatives. *In vitro* and *in vivo* observations were mostly consistent with some notable exceptions. For example, the MEG-3 C-terminus was more efficient at condensation than the MEG-3 IDR *in vivo*. This difference could also be observed *in vitro* when MEG-3 condensation was assayed in the presence of PGL-3, which solubilizes the MEG-3 IDR (Figure 2c). When MEG-3 was assayed alone, however, a difference between the MEG-3 c-terminus and IDR condensation could only be seen at the lowest concentration tested (Figure 2A). A second difference between *in vitro* and *in vivo* assays was observed when examining the HMGL motif. Mutations in this domain greatly disrupted MEG/PGL co-assembly *in vivo* but had no apparent effect in the *in vitro* co-assembly assay. Mutations in the HMGL domain did, however, lower MEG-3’s affinity for the PGL proteins when assayed by GST pull down. The GST pull down assay is done under more stringent conditions (SDS and high salt) than the condensation assays and may therefore be better suited to reveal affinity differences sufficient to disrupt protein interactions in the crowded cellular milieu. We conclude that, while *in vitro* experiments are excellent tools to reveal self-assembly principles, condensation assays can lead to conclusions (e.g. HMGL is dispensable for MEG-3/PGL co-assembly) that do not necessarily hold *in vivo*.

### Assembly of MEG-3/PGL co-condensates depends on interactions between MEG and PGL proteins and does not require RNA

The MEG-3 c-terminus is a 318 aa sequence with regions of predicted low disorder including an HMG-like motif. We have found that the MEG-3 c-terminus is sufficient to form condensates that dock on PGL droplets *in vitro* and *in vivo*. Docking of P bodies on stress granules has been proposed to involve RNA:RNA duplexes (Tauber et al., 2020). In contrast, we find that docking of MEG-3 condensates on PGL condensates does not require RNA *in vitro* and can occur in the absence of any visible RNA enrichment *in vivo*. Mutations in the HMGL domain prevent the association of MEG-3 and PGL condensates *in vivo* and lower the affinity of the MEG-3 c-terminus PGL proteins *in vitro*. Together, these observations strongly suggest that the association between MEG-3 and PGL condensates depends primarily on protein-protein interactions.

We previously showed that PGL-3 condensates stimulate MEG-3 condensation *in vitro* and *in vivo* (Wang et al., 2014, Putnam et al., 2019). consistent with this, we report here that mutations in the HMGL domain that prevent MEG-3/PGL co-assembly also reduce MEG-3 condensation efficiency *in vivo*. An attractive possibility is that high affinity MEG-3/PGL binding, mediated in part by the HMGL motif, recruits MEG-3 molecules from the cytoplasm to the surface of PGL droplets, driving their condensation. We note that PGL molecules are unlikely to be the only binding partners that stimulate MEG condensation, since *pgl-1;pgl-3* mutants embryos still assemble MEG granules that support RNA assembly (although not as robustly as wild-type; Lee et al., 2020).

### Efficient MEG-3/PGL co-assembly correlates with stabilization of PGL droplets in germ plasm

We previously reported that enrichment of PGL droplets to the posterior of the zygote requires *meg-3* (Smith et al., 2016; Wang et al., 2014). Our new findings suggest that this activity is linked to MEG-3’s ability to associate stably with the PGL interface. MEG-3_Cterm_, which is sufficient for MEG-3/PGL co-assembly, is sufficient to localize PGL in zygotes. conversely, MEG-3_HMGL_ condensates, which do not interact stably with PGL-3 condensates, fail to enrich PGL condensates in the posterior. PGL localization involves preferential growth and dissolution of PGL droplets in the anterior and posterior, respectively. One possibility is that tight binding of MEG condensates lowers the surface tension of PGL droplets allowing MEG/PGL coassemblies in the posterior to grow at the expense of the less stable, “naked” PGL droplets in the anterior.

What enriches MEG-3 condensates in the posterior? We previously hypothesized that MEG-3 asymmetry is driven by a competition for RNA between the MEG-3 IDR and MEX-5, an RNA-binding protein that acts as an RNA sink in the anterior (Smith et al., 2016). Our finding that condensation of MEG-3_Cterm_ is restricted to the zygote posterior despite uniform distribution in the cytoplasm suggest that additional mechanisms acting on the MEG-3 C-terminus contribute to MEG-3 regulation in space. Consistent with this view, a recent study examining MEG-3 dynamics by single-molecule imaging (Wu et al., 2019) found that the slowly-diffusing MEG-3 molecules that populate the MEG-3 gradient in the cytoplasm represent a distinct population of MEG-3 molecules from those that associate with PGL droplets. Although the MEG-3 cytoplasmic gradient does not contribute to P granule asymmetry directly, it may serve to maintain high levels of MEG-3 in P blastomeres to sustain PGL asymmetry through the P_4_ stage. Consistent with this view, the MEG-3_Cterm_, which does not enrich in a gradient, is not sufficient to localize PGL in P blastomeres past the 4-cell stage.

### MEG-3 condensation on PGL droplets creates a platform for RNA recruitment

The MEG-3 IDR binds RNA with high affinity *in vitro* but is not sufficient to enrich RNA *in vivo* despite forming condensates. RNA recruitment also requires the MEG-3 C-terminus including the HMG-like motif. These observations suggest that condensation of the MEG-3 C terminus is essential to build a protein scaffold that can support RNA recruitment *in vivo*. Separate domains for RNA binding and protein condensation have also been observed for other germ granule scaffolds. For example, the Balbiani body protein Xvelo uses a prion-like domain to aggregate and a separate RNA-binding domain to recruit RNA (Boke et al., 2016). Similarly, condensation of Drosophila Oskar does not require the predicted Oskar RNA-binding domain, although this domain augments condensation (Kistler et al., 2018). These observations parallel our findings with MEG-3 and contrast with recent findings reported for the stress granule scaffold G3BP. Condensation of G3BP *in vitro* requires RNA and two C-terminal RNA-binding domains. A N-terminal dimerization domain is also required but, unlike the prion-like domain of Xvelo or the C-terminus of MEG-3, is not sufficient to drive condensation on its own. Dimerization of G3BP is thought to enhance LLPS indirectly by augmenting the RNA-binding valency of G3BP complexes. G3BP also contains an inhibitory domain that gates its RNA-binding activity and condensation at low RNA concentrations. This modular organization ensures that G3BP functions as a sensitive switch that initiates LLPS when sufficient RNA molecules are available to cross-link G3BP dimers into a large network (Guillén-Boixet et al., 2020; Yang et al., 2020). Stress granules are transient structures that form under conditions of general translational arrest where thousands of transcripts are released from ribosomes. In contrast, germ granules are long-lived structures that assemble in translationally-active cytoplasm and recruit only a few hundred specific transcripts (~500 in *C. elegans* embryos) (Jamieson-Lucy and Mullins, 2019; Lee et al., 2020; Trcek and Lehmann, 2019; Updike and Strome, 2010). One possibility is that protein-based condensation mechanisms may be better suited to assemble long-lived granules able to capture and retain rare transcripts. By concentrating IDRs with affinity for RNA, protein scaffolds could act as seeds for localized LLPS to amplify protein and RNA condensation. consistent with this view, IDRs have been observed to undergo spontaneous LLPS in cells when artificially tethered to protein modules that self-assemble into large multimeric structures (Nakamura et al., 2019). A challenge for the future will be to understand the mechanisms that regulate the assembly and disassembly of protein scaffolds at the core of germ granules.

## METHODS

### Worm handling, maternal-effect sterility counts

*C. elegans* was cultured at 20° C according to standard methods (Brenner, 1974). To measure maternal-effect sterility, ten gravid adults were picked to an OP50 plate and allowed to lay eggs for ~2 hours, then removed. Adult progeny were scored for empty uteri (white sterile phenotype) under a dissecting microscope.

### Identification of MEG-3 HMG-like region

MEG-3 and MEG-4 protein sequences were aligned with HMG boxes from GCNA proteins of *Caenorhabditis* and example vertebrates along with the canonical HMG box of mouse SOX3 using MUSCLE (Edgar, 2004). Alignment was manually adjusted according to the published CGNA HMG Hidden Markov Model (Carmell et al., 2016). Amino acids were chosen for mutation based on conservation in nematodes.

### CRISPR genome editing

Genome editing was performed in *C. elegans* using CRISPR/Cas9 as described in (Paix et al., 2017). Strains used in this study along with guides and repair templates are listed in Supplementary Table 1. Some strains were generated in two steps. For example, MEG-3_HMGL-_ was generated by deleting the entire HMGL-like motif in a first step (JH3632), and inserting a modified HMG-like motif with the desired mutations in a second step (JH3861). Genome alterations were confirmed by Sanger sequencing and expression of tagged strains was verified by immunostaining and western blotting (Sup. Figure 3B).

### Statistical Analysis and plotting

On all scatterplots, central bars indicate the mean and error bars indicate one standard deviation. Unless otherwise indicated, differences within three or more groups were evaluated using a one-factor ANOVA and differences between two groups using an unpaired Student’s t-test.

### Confocal Imaging

Fluorescence confocal microscopy for figure 2C, supplementary figures 2A, 3B and 4A performed using a Zeiss Axio Imager with a Yokogawa spinning-disc confocal scanner. Fluorescence confocal microscopy for all other figures was performed using a custom built inverted Zeiss Axio Observer with CSU-W1 Sora spinning disk scan head (Yokogawa), 1X/2.8x relay lens (Yokogawa), fast piezo z-drive (Applied Scientific Instrumentation), and a iXon Life 888 EMCCD camera (Andor). Samples were illuminated with 405/488/561/637nm solid-state laser (Coherent), using a 405/488/561/640 transmitting dichroic (Semrock) and 624-40/692-40/525-30/445-45nm bandpass filter (Semrock) respectively. Images from either microscope were taken with using Slidebook v6.0 software (Intelligent Imaging Innovations) using a 40x-1.3NA/63X-1.4NA objective (Zeiss) depending on sample.

### Immunostaining

**Supplementary Table 1:**
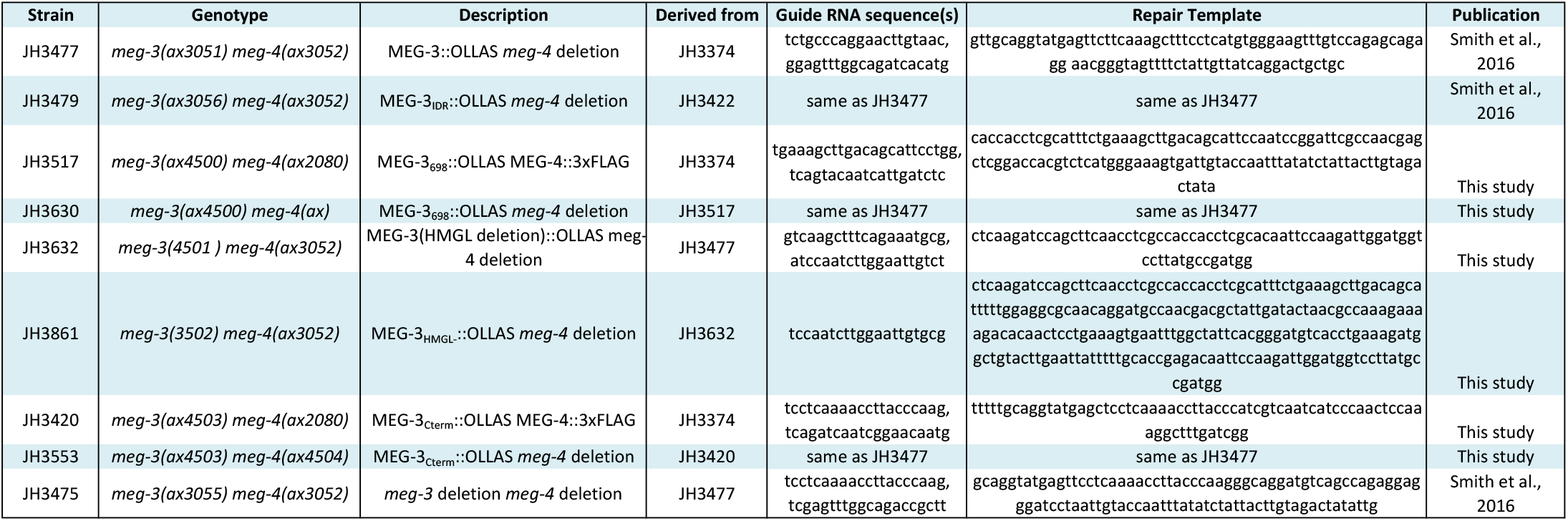
*C. elegans* strains used in this study, generated by CRISPR/Cas9 genome editing

Adult worms were placed into M9 on poly-l-lysine (0.01%) coated slides and squashed with a coverslip to extrude embryos. Slides were frozen by laying on aluminum blocks pre-chilled with dry ice for >5 min. Embryos were permeabilized by freeze-cracking (removal of coverslips from slides) followed by incubation in methanol at −20°C for >15 min, and in acetone −20°C for 10 min. Slides were blocked in PBS-Tween (0.1%) BSA (0.5%) for 30min at room temperature, and incubated with 50 ul primary antibody overnight at 4°C in a humid chamber. For co-staining experiments, antibodies were applied sequentially (OLLAS before KT3, K76) to avoid cross reaction. Antibody dilutions (in PBST/BSA): KT3 (1:10, DSHB), K76 (1:10 DSHB), Rat αOLLAS-L2 (1:200, Novus Biological Littleton, CO), Secondary antibodies were applied for 2 hr at room temperature. Samples were mounted Prolong Diamond Antifade Mountant or VECTASHIELD Antifade Mounting Media with DAPI. Embryos were staged using DAPI stained nuclei and 25 confocal slices spaced 0.18 microns apart and centered on the P cell nucleus were taken using a 63x objective. Unless otherwise indicated, images presented in figures are maximum projections.

### Quantification of immunostaining images

All analysis was performed in ImageJ. For measurements of embryos/cells, confocal stacks were sum projected and the integrated density was measured within a region of interest. For measurements of condensate intensity, the 3D objects counter function was used on the full confocal stack confined to a region of interest drawn around the P cell and including objects on edges. The integrated density for all identified particles was summed to give the total intensity in condensates.

### Single molecule fluorescence in situ hybridization (smFISH)

smFISH probes were designed using Biosearch Technologies’s Stellaris Probe Designer, with the fluorophor Quasar670. For sample preparation, embryos were extruded from adults on poly-l-lysine (0.01%) slides and subjected to freeze-crack followed by methanol fixation at −□20° C for >15minutes. Samples were washed five times in PBS-Tween (0.1%) and fixed in 4% PFA (Electron Microscopy Science, No.15714) in PBS for one hour at room temperature. Samples were again washed four times in PBS-Tween (0.1%), twice in 2x SCC, and once in wash buffer (10% formamide, 2x SCC) before blocking in hybridization buffer (10% formamide, 2x SCC, 200 ug/mL BSA, 2 mM Ribonucleoside Vanadyl Complex, 0.2 mg/mL yeast total RNA, 10% dextran sulfate) for >30 min at 37° C. Hybridization was then conducted by incubating samples with 50nM probe solutions in hybridization buffer overnight at 37° C in a humid chamber. Following hybridization, samples were washed twice in wash buffer at 37° C, twice in 2x SCC, once in PBS-Tween (0.1%) and twice in PBS. Samples were mounted Prolong Diamond Antifade Mountant.

### Quantification of *in situ* hybridization images

All measurements were performed on a single confocal slice centered on the P cell nucleus in ImageJ. For early embryos where there is distinct punctate signal (1 and 4 cell stage Figure 5B,C, Figure 5 – figure 1 B), a region of interest was drawn, the Analyze Particles feature was used with a manual threshold to identify and measure the integrated density of the puncta. The raw integrated density for all particles in the region of interest was summed to give the total intensity of the mRNA in that region. For 28 cell embryos (Fig, 5D) a region of interest was drawn around the P_4_ blastomere and the intensity of that region was divided by the intensity of a region of the same size in the anterior soma.

### Western blotting of embryonic lysates

Worms were synchronized by bleaching to collect embryos, shaken approximately 20hrs in M9, then plating on large enriched peptone plates with a lawn of *E. Coli* NA22 bacteria. Embryos were harvested from young adults (66 hours after starved L1 plating) and sonicated in 2% SDS, 65 mM Tris pH 7, 10% glycerol with protease and phosphatase inhibitors. Lysates were spun at 14,000 rpm for 30 min at 4° c and cleared supernatants were transferred to fresh tubes. Lysates were run on 4-12% Bis-Tris pre-cast gels (Bio-Rad Hercules, cA). Western blot transfer was performed for 1 hr at 4°c onto PVDF membranes. Membranes were blocked overnight and washed in 5% milk, 0.1% Tween-20 in PBS; primary antibodies were incubated overnight at 4° c; secondary antibodies were incubated for two hours at room temperature. Membranes were first probed for OLLAS then stripped by incubating in 62.5mM Tris Hcl pH6.8, 2% SDS, 100mM ß-mercaptoethanol at 42° c. Membranes were then washed, blocked and probed for α-tubulin. Antibody dilutions in 5% milk/PBST: Rat α OLLAS-L2 (1:1000, Novus Biological Littleton, cO), Mouse α tubulin (1:1000, Sigma St. Louis, MO).

### His-tagged protein expression, purification and labeling

#### Expression and purification of MEG-3 His-tagged fusion proteins

MEG-3 full-length (aa1-862), IDR (aa1-544), cterm (aa545-862), and HMGL-proteins were fused to an N-terminal 6XHis tag in pET28a and expressed and purified from inclusion bodies using a denaturing protocol (Lee et al., 2020)

Purification of MBP-TEV-PGL-3 was expressed and purified as described (Putnam et al., 2019) with the following modifications: MBP was cleaved using homemade TEV protease instead of commercial. A plasmid expressing 8X-His-TEV-8X-Arg tag protease was obtained from Addgene and purified according to the published protocol (Tropea et al., 2009). Before loading cleaved PGL-3 protein on to a heparin affinity matrix, cleaved MBP-6X-His and 6X-His-TEV protease were removed using a HisTRAP column (GE Healthcare).

#### Protein labeling

Proteins were labeled with succinimidyl ester reactive fluorophores from Molecular Probes (Alexa Fluor™ 647 or DyLight™ 488 NHS Ester) following manufacturer instructions. Free fluorophore was eliminated by passage through three Zeba™ Spin Desalting columns (7K MWcO, 0.5 mL) into protein storage buffer. The concentration of fluorophore-labeled protein was determined using fluorophore extinction coefficients measured on a Nanodrop ND-1000 spectrophotometer. Labeling reactions resulted in ~ 0.25-1 label per protein. Aliquots were snap frozen and stored. In phase separation experiments, fluorophore-labeled protein was mixed with unlabeled protein for final reaction concentrations of 25-100 nM of fluorophore labeled protein.

### *In vitro* transcription and labeling of RNA

mRNAs were transcribed using T7 mMessageMachine (Thermofisher) using manufacturer’s recommendation as described (Lee et al., 2020). Template DNA for transcription reactions was obtained by PCR amplification from plasmids. Free NTPs and protein were removed by lithium chloride precipitation. RNAs were resuspended in water and stored at −20°C. The integrity of RNA products was verified by agarose gel electrophoresis.

### *In vitro* condensation experiments and analysis

Protein condensation was induced by diluting proteins out of storage buffer into condensation buffer containing 25 mM HEPES (pH 7.4), salt adjusted to a final concentration of 150 mM (37.5 mM KCl, 112.5 mM NaCl), and RNA. For MEG-3 and PGL-3 co-condensate experiments with RNA, we used 150 nM MEG-3, 1.8 μM PGL-3 and 20 ng/μL RNA. For co-assembly experiments in the absence of RNA, we used 150 nM MEG-3, 5 μM PGL-3. MEG-3 and PGL-3 solutions contained 25 nM fluorescent trace labels with either 488 or 647 (indicated in figure legends). MEG-3 condensation reactions were incubated at room temperature for 30 min before spotting onto a No. 1.5 glass bottom dish (Mattek) and imaged using a 40x oil objective (Figure 2A). MEG-3 and PGL-3 co-condensate were imaged using thin chambered glass slides (Erie Scientific Company 30-2066A) with a coverslip (Figure 2C). Images are single planes acquired using a 40x oil objective over an area spanning 171 x 171 μm.

To quantify the relative intensity of MEG-3 in condensates, a mask was created by thresholding images, filtering out objects of less than 4 pixels to minimize noise, applying a watershed filter to improve separation of objects close in proximity, and converting to a binary image by the Otsu method using the nucleus counter cookbook plugin. Minimum thresholds were set to the mean intensity of the background signal of the image plus 1-2 standard deviations. The maximum threshold was calculated by adding 3-4 times the standard deviation of the background. Using generated masks, the integrated intensity within each object was calculated. To remove non-specific background signal the mean intensity of an image field in the absence of the labeled component was subtracted from each pixel yielding the total intensity of each object. The relative intensity in each condensate was normalized to the mean intensity of WT MEG-3. Each data point represents the average of 3 experimental replicates. Each replicate contained 4 images each spanning an area of 316.95 x 316.95 μm.

### RNA binding by fluorescence filter binding

Proteins were step-dialyzed from 6M Urea into 4.5M Urea, 3M Urea, 1.5M Urea, and 0M Urea in MEG-3 Storage Buffer (25 mM HEPES, pH 7.4, 1M Nacl, 6 mM ß-mercaptoethanol, 10% Glycerol). RNA binding reactions consisted of 50 nM 3’ Fluorescein-labeled 30U RNA oligonucleotides incubated with protein for 30 min at room temperature (final reaction conditions 3.75 mM HEPES, 150mM Nacl, 0.9mM, 0.9mM ß-mercaptoethanol, 1.5% glycerol, 10mM Tris Hcl). Fluorescence filter binding protocol was adapted from a similar protocol using radiolabeled RNA(Rio, 2012). Briefly, a pre-wet nitrocellulose was placed on top of Hybond-N+ membrane in a dot-blot apparatus, reactions were applied to the membranes, then washed 2x with 10mM Tris Hcl. Membranes were briefly dried in air, then imaged using a typhoon FLA-9500 with blue laser at 473 nm. Fraction of RNA bound for each reaction was calculated by dividing the fluorescence signal on the nitrocellulose membrane by the total signal from both membranes. K_d_ was calculated by plotting the bound fraction of RNA as a function of protein concentration and fitting to the following equation in Prism8 where P is the protein concentration in nM, B is the bound fraction of RNA with non-specific binding subtracted, B_max_ is the maximum specific binding, K_d_ is the concentration needed to achieve a half-maximum binding at equilibrium, and h is the Hill slope.

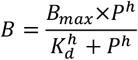

### GST pull-downs

GST fusion proteins were cloned into pGEX6P1 (GE Healthcare, Pittsburgh, PA). MBP fusion proteins were cloned into pJP1.09, a Gateway-compatible pMAL-c2x (Pellettieri et al., 2003). Proteins were expressed in Rosetta™ *E. coli* BL21 cells grown for approximately four hours at 37° c then induced with 1mM IPTG and grown overnight at 16° c. 200 mg of bacterial pellet of GST fusion proteins was resuspended in 50mM HEPES, 1 mM EGTA, 1 mM Mgcl_2_, 500 mM Kcl, 0.05% NP40, 10% glycerol, pH 7.4 with protease and phosphatase inhibitors, lysed by sonication, and bound to magnetic GST beads. Beads were washed and incubated with MBP fusion proteins at 4°c for 1 hr in the same buffer as for lysis. After washing, beads were eluted by boiling and eluates were loaded on SDS-PAGE. Western blot transfer was performed for 1 hr at 4°c onto PVDF membranes. Membranes were blocked and washed in 5% milk, 0.1% Tween-20 in PBS and incubated with HRP conjugate antibodies. Antibody dilutions in 5% milk/PBST: anti-MBP HRP conjugated, 1:50,000 (NEB, and anti-GST HRP conjugates, 1:2,000 (GE Healthcare,). Scanned western blot films were quantified using the gel analysis tool in ImageJ.

## ACKNOWLEDGEMENTS

We thank the Johns Hopkins Neuroscience Research Multiphoton Imaging core (NS050274) and the Johns Hopkins Integrated Imaging center (S10OD023548) for excellent microscopy support. We thank the Page lab for assistance identifying the HMG-like motif, the Nathans lab for the OLLAS antibody, and Addgene for TEV protease from the Waugh lab. We thank Deepika calidas for assistance in strain construction, Baltimore Worm club and the Seydoux lab for many helpful discussions. This work was supported by the National Institutes of Health (Grant number 5R37HD037047). GS is an investigator of the Howard Hughes Inistitute.

---

**Figure 2 - figure supplement 1.**
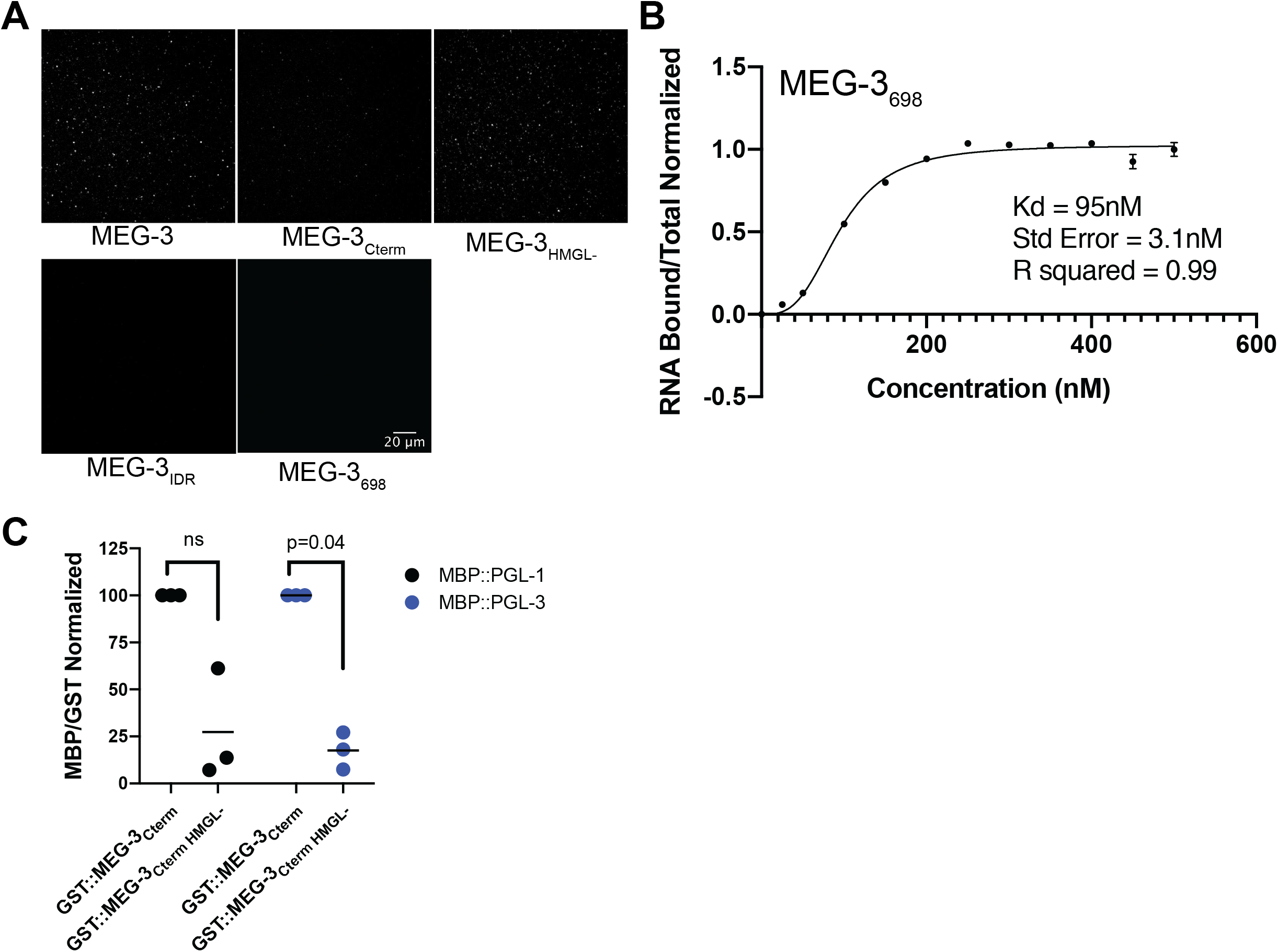
**(A)** Representative photomicrographs of Alexa488 trace-labeled MEG-3 and MEG-3 derivatives in condensation buffer with 20 ng/μL *Y51F10.2* mRNA. **(B)** RNA binding curve for MEG-3_698_. Protein concentration is plotted on the X-axis. The ratio of bound RNA to total RNA, normalized to the ratio at the maximum concentration is plotted on the Y-axis. Dots indicate the mean of three replicates and the error bars the standard deviation. Curve fit and Kd calculation based on specific binding with Hill slope (Methods) **(C)** Scatterplot showing the ratio of the indicated MBP fusions to GST:MEG-3_Cterm HMGL-_ normalized to the ratio of the same MBP fusion to GST:MEG-3_Cterm_ from the same experiment. Each dot represents an independent pull-down experiment. P values indicated above were calculated by an paired ratio t-test of the GST:MEG-3_Cterm_ and GST:MEG-3_Cterm HMGL-_ ratios before normalization.

**Figure 3 - figure supplement 1.**
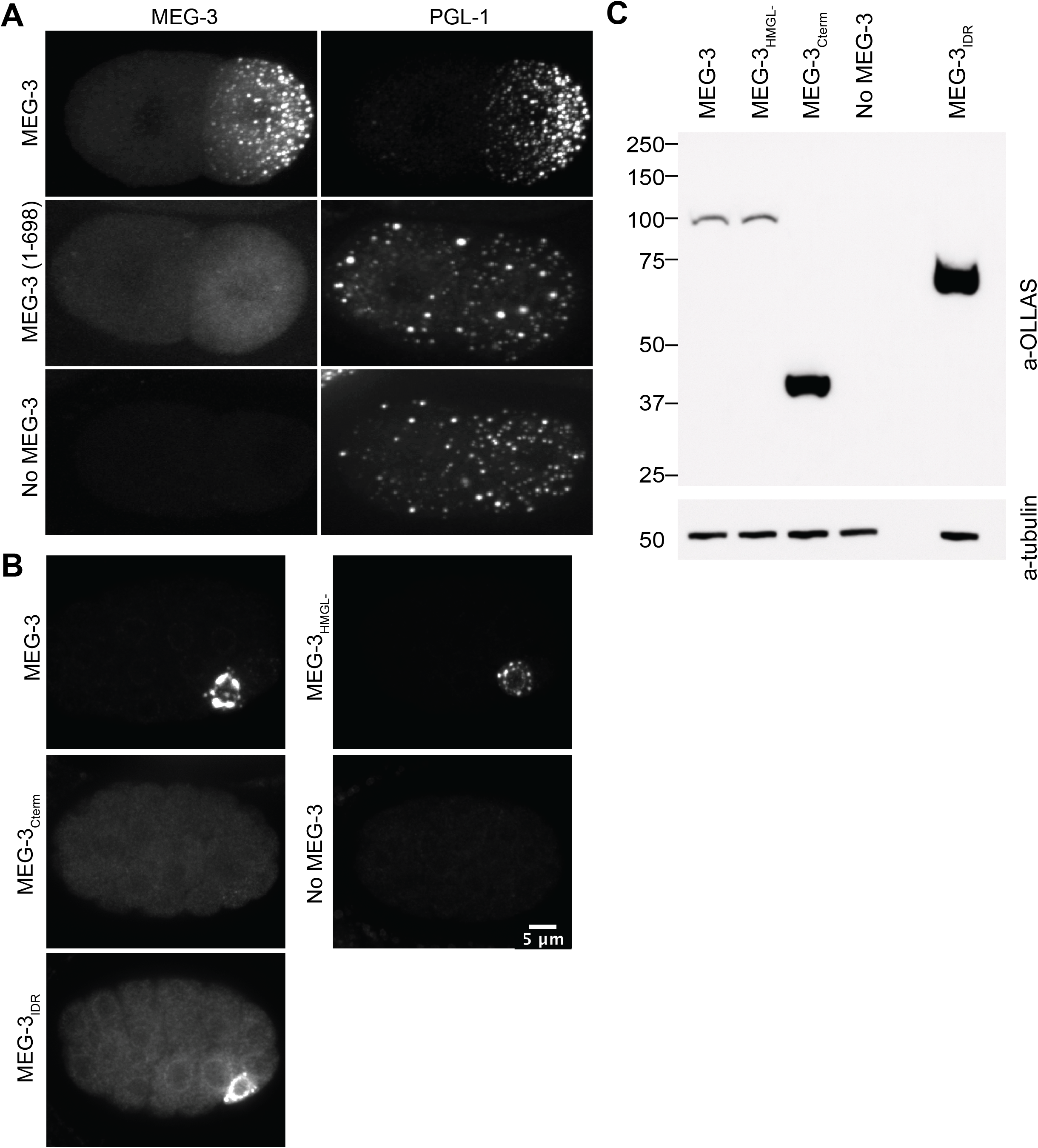
**(A)** Representative photomicrographs of two-cell embryos expressing the indicated MEG-3 derivatives and immunostained for MEG-3 (anti-OLLAS antibody) and PGL-1 (anti-PGL-1 antibody). **(B)** Representative photomicrographs of sum projections of 28-cell stage embryos expressing the indicated MEG-3 derivatives and immunostained for MEG-3. **(C)** Westerns of mixed-stage embryos (1-100 cell stage) harvested from synchronized worms expressing the indicated OLLAS-tagged MEG-3 derivatives.

**Figure 4 - figure supplement 1.**
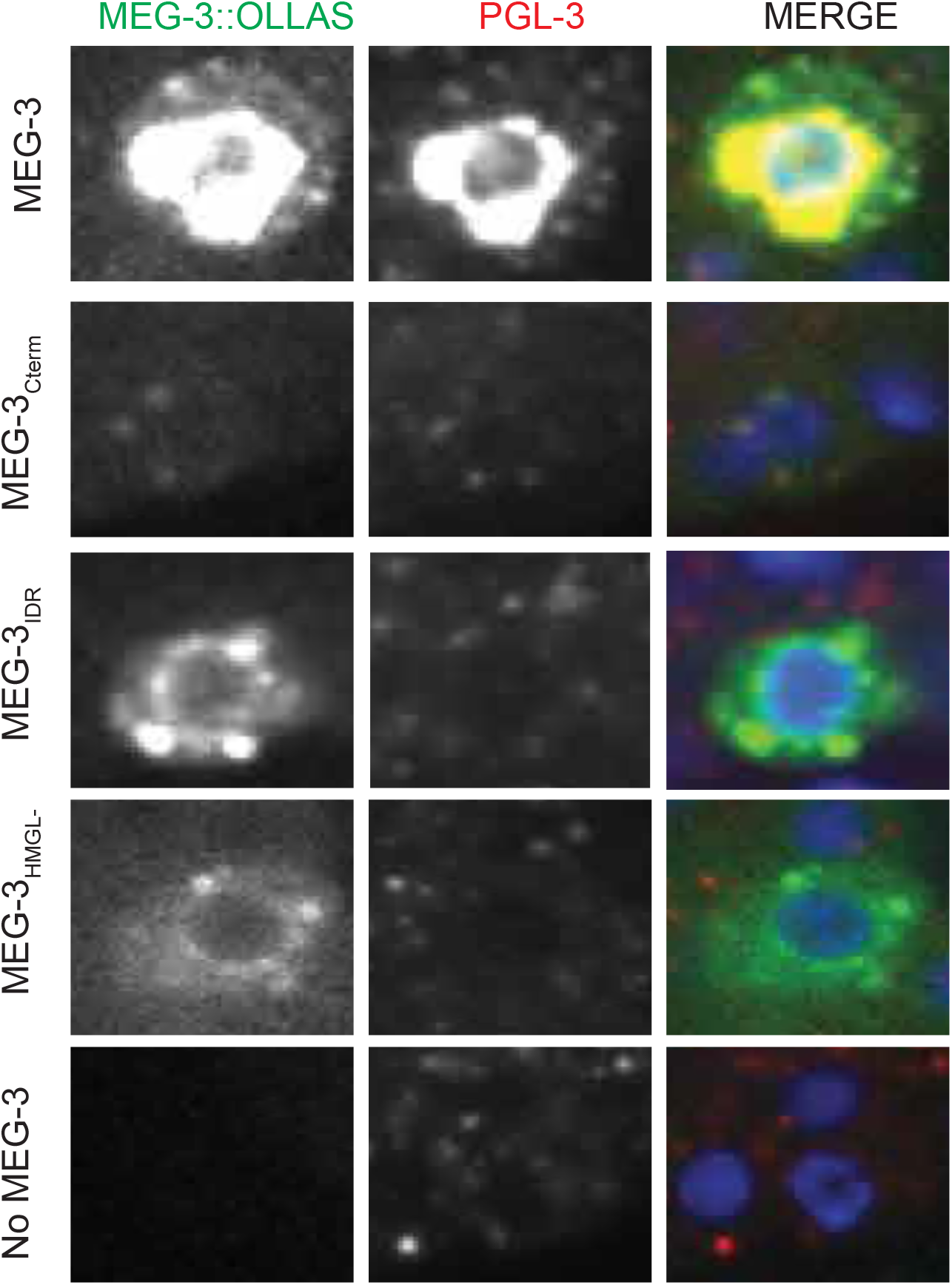
Representative photomicrographs of a single confocal slice centered on the P_4_ blastomere of embryos expressing the indicated MEG-3 mutants and immunostained for MEG-3 (anti-OLLAS antibody) and PGL-3 (anti-PGL-3 antibody). Note colocalization of MEG-3_Cterm_ and PGL-3. MEG-3_Cterm_ is present at lower level in P_4_ compared to other MEG-3 derivatives, consistent with lack of enrichment in germ plasm starting in the 1-cell stage (Figure 3).

**Figure 5 - figure supplement 1.**
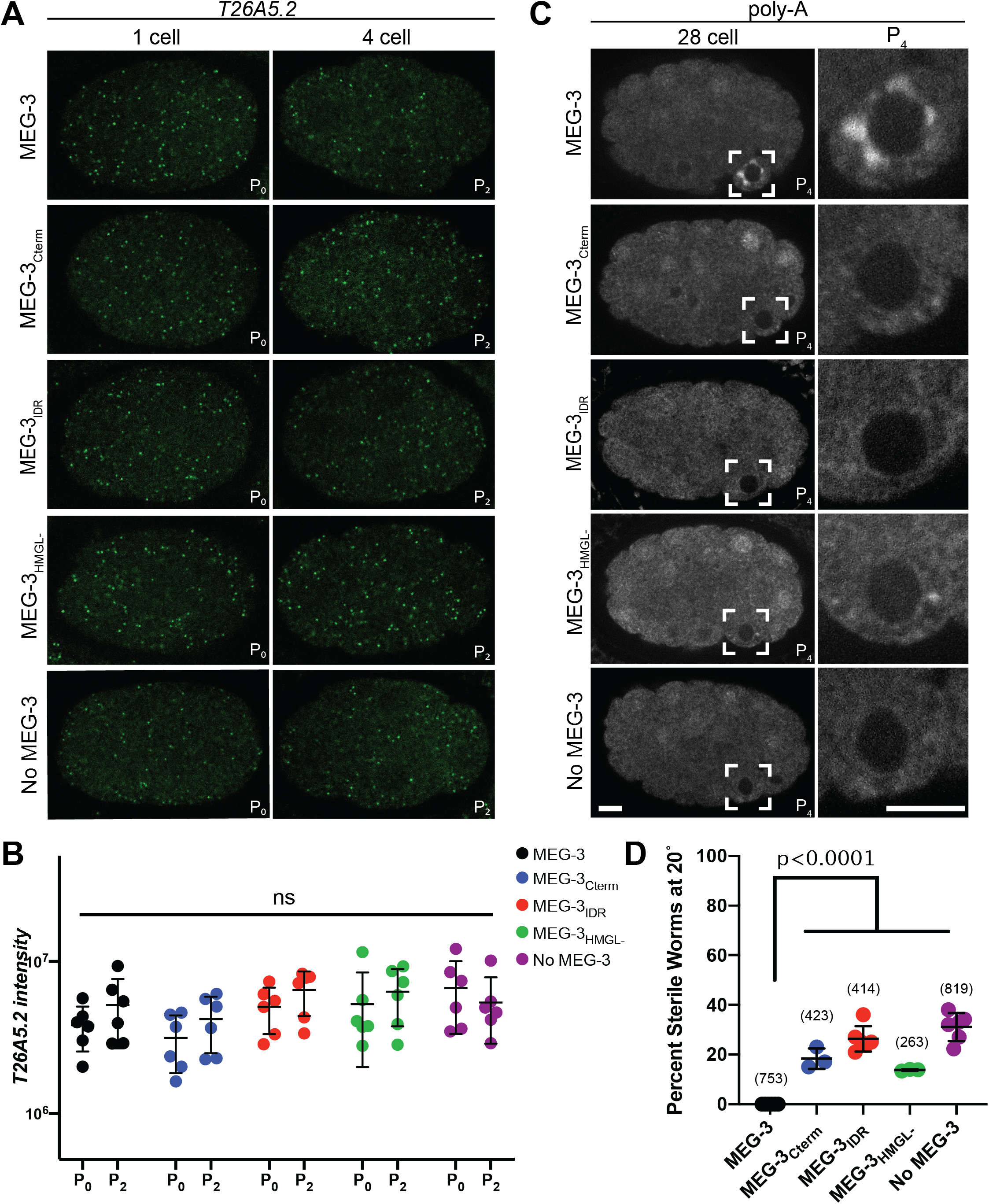
**(A)** Representative photomicrographs of single confocal slices of fixed embryos expressing the indicated MEG-3 variants and hybridized to fluorescent probes complementary to the mRNA *T26A5.2*. Images are of the same embryo and confocal slice as Figure 5A. **(B)** Scatterplot of the intensity the *T26A5.2* mRNA signal in PO and P_2_ embryos expressing the indicated MEG-3 derivatives. Each dot represents an embryo analysed in Fig 5B. **(C)** Representative photomicrographs of single confocal slices of fixed embryos expressing the indicated MEG-3 variants and hybridized to oligo-dT fluorescent probes to detect polyadenylated mRNAs. **(D)** Plot of the percentage of sterile worms from mothers expressing the indicated MEG-3 derivatives raised at 20°. Each dot represents the 2-hour brood of 10 mothers (Methods). Total number of worms scored is shown above.

